# Chemical Profiling of Antimicrobial Metabolites from Halophilic Actinomycete *Nocardiopsis_sp. Al-H10-1* (KF384482) Isolated from Alang, Gulf of Khambhat, India

**DOI:** 10.1101/2021.06.12.448169

**Authors:** Nisha Trivedi, Jignasha Thumar

## Abstract

The overuse of antibiotics has resulted in the development of drug resistant, a major problem in disease curing processes i.e. development of drug resistance. The World Health Organization (WHO) released its first list of the most concerning pathogens for human health in 2017 which suggested that there are total 12 bacterial families which have developed multiple drug resistance and for which novel antibiotics are required immediately (WHO 2017). There is a requirement to explore some novel compounds to overcome this issue. Thus our study aimed at exploration of marine actinomycetes as a valuable resource for novel products with antimicrobial properties. The halophilic actinomycete *Nocardiopsis_sp. Al-H10-1* (KF384482) was isolated from saline water (20 m away from shore) of Alang coast (Gulf of Khambhat), Bhavnagar, Gujarat, India. The isolate Al-H10-1 was identified as *Nocardiopsis sp*. through rigorous morphological and cultural characteristics; the species was confirmed through 16s rRNA phylogenetic analysis. The antimicrobial potential of *Nocardiopsis sp. Al-H10-1* was assessed against a range of Gram-positive and Gram-negative bacteria as well as three fungi, there it demonstrated antimicrobial activity against four Gram negative bacteria and one Gram positive bacteria. Further active antimicrobial compounds present in ethyl acetate extract was identified using Gas Chromatography-Mass Spectroscopy (GC-MS). GC-MS analysis showed the presence of 17 compounds which included antimicrobial compounds like 2, 4-bis (1, 1-dimethylethyl)-Phenol, Dibutyl phthalate as well as various types of alkanes and their derivatives.

## INTRODUCTION

The World Health Organization (WHO) released its first list of the most concerning pathogens for human health in 2017 which suggested that there are total 12 bacterial families who have developed multiple drug resistance and for which novel antibiotics are required immediately (WHO, 2017). The study by O ‘Neill suggests that every year approximately 7,00,000 people die because of drug resistant infections globally and if current trends continue, it will be increased by 10 million people per year by 2050 (O ‘Neill J, 2016). The multiple drug resistance can be developed in organisms by various types of mechanisms such as presence of antibiotic degrading enzymes, antibiotic altering enzymes, gene transfer processes like conjugation, transformation, transduction etc. So there is a need for searching naturally occurring novel antimicrobial compounds.

The natural products and microbial metabolites are used as the main bioactive scaffold for the development of the novel antibiotics instead of using the large synthetic combinatorial libraries of molecules to develop novel drugs (Challinor VL and Bode HB, 2015). During 1981 to 2014, total 1211 small molecule drugs were approved by the United States Food and Drug Administration (FDA), from which approximately 65% was accounted for chemicals produced in nature, or compounds based on them (Newman DJ and Cragg GM, 2016). Actinomycetes have always been known as the biotechnologically and industrially important group of microorganisms as they are the great producers of many primary and secondary metabolites useful in a variety of fields including healthcare, agriculture, veterinary etc. (Barka EA et. al., 2016). Various unexplored or underexplored ecosystems are the most promising source of novel actinomycetes able to produce large numbers of secondary metabolites (Dhakal D et. al., 2016; Subramani R and Aalbersberg W, 2013).

The Gulf of Khambhat is a coastal region situated in the Bhavnagar District of the Gujarat State of India. The uniqueness of this region is, its salinity and alkalinity which harbors various unidentified, unique haloalkaliphilic bacterial species which can produce valuable secondary metabolites. Alang is a small town located on the Gulf of Khambhat, considered as the world ‘s largest graveyard of ships. It is well known for its ship breaking and recycling activities, because of which it is also known as the most polluted coastal region (Patel AB et. al., 2018).

## MATERIALS AND METHODS

### Isolation

The halophilic actinomycete, *Nocardiopsis_sp. Al-H10-1* was isolated from saline water (20 m away from shore) of Alang coast (Gulf of Khambhat), Bhavnagar, Gujarat, India. The water sample was filtered with whatman filter paper to remove fine soil particles and the filtrate was serially diluted. From the highest dilution containing tube, 0.1 ml of the sample was spreaded separately on starch agar, starch casein agar and actinomycetes isolation agar (Hi-Media, India), containing 5-10% w/v NaCl. The plates were incubated at 30°C; on sixth day a typical chalky white colony was picked up and re-streaked on starch agar slants supplemented with 10% w/v NaCl to ensure the purity of the colony (Chakrabarti T, 1998). The culture was maintained at 4°C.

### Morphological and Biochemical Characteristics and Antibiotic Susceptibility Test

Morphological, physiological and biochemical characteristics of the isolate were determined using standard methods. The morphological characteristics were determined by growing the isolate on a starch agar plate (with 10% NaCl). After seven days of incubation, the colony characteristics were noted and Gram ‘s staining was performed. Basic biochemical tests were performed for primary characterization of the isolate based on Bergey ‘s manual of systematic Bacteriology Vol. IV (Garrity GM et. al., 2004), which included including carbon sources utilization (Mannitol, Sucrose, Xylose, Glucose, Maltose, and Lactose), Methyl red test, Voges-Proskauer test, H_2_S production, Catalase test, Indole production, Citrate utilization, Starch hydrolysis, Casein hydrolysis, Gelatin hydrolysis, Nitrate reduction, Urea hydrolysis, Ammonia production, and Triple sugar iron test. All media and test reagents were prepared as mentioned by Cappuccino JG and Sherman N (2004). Antibiotic susceptibility and resistance pattern of the isolate were checked by disc diffusion method (Kumar PS et. al., 2014). The test was conducted against total 13 antibiotics using Starch agar plates with 10% NaCl.

### Molecular Identification of Isolate by 16S rRNA

The molecular identification of the isolate Al-H10-1 was carried out by 16S rRNA gene sequencing. The 16S rRNA gene was amplified using universal primers 518F 5 ‘ccagcagccgcggtaatacg3 ‘ and 800R 5 ‘taccagggtatctaatc3 ‘. PCR products were purified and sequenced. The resultant sequences aligned within the NCBI database (National Centre for Biotechnology Information) using BLASTN. The phylogenetic tree was constructed using neighbor-joining with Kimura 2-state parameter and pairwise-deletion model analysis implemented in the program MEGA software version X (Kumar S et. al., 2018) and also evaluated further in a bootstrap analysis of 1,000 replicates. The 16S rRNA gene sequence of the isolate has been submitted to NCBI, GenBank, Maryland, USA.

### The Antimicrobial Potential

Antimicrobial activity of *Nocardiopsis_sp. Al-H10-1* was checked by spot inoculation method (Kumar N et. al., 2010) using starch agar (10 % w/v NaCl). The spore suspension of the isolate was spotted on the medium and incubated at 28°C until sporulation. The test organisms, procured from MTCC, Chandigarh (ATCC equivalent), were used to check antimicrobial activity. These included Gram-positive organisms: *Bacillus subtilis, Staphylococcus aureus, Micrococcus luteus, Staphylococcus epidermidis, Bacillus megaterium, Bacillus cereus;* Gram-negative organisms: *Pseudomonas aeruginosa, Escherichia coli, Enterobacter aerogenes, Serratia marcescens, Shigella flexneri, Salmonella enteric typhimurium, Proteus vulgaris, Klebsiella pneumonia* and fungi: *Aspergillus niger, Fusarium oxysporum, Candida albicans*. The test organisms were grown in nutrient broth at 37°C for 24 hours. The molten nutrient agar, with 1% activated test culture, was poured on the sporulated *Nocardiopsis_sp. Al-H10-1*. After the incubation of 24 hours at 37°C, the zone of inhibition was measured for each test organism.

### Optimization of Growth Conditions for Antimicrobial Compound Production

To achieve the highest production of antimicrobial compounds, effect of various growth conditions were studied by the one variable at a time (OVAT) method. To find out the optimized medium for the growth of the organism as well as for antimicrobial compounds production, the spore suspension of *Nocardiopsis_sp. Al-H10-1* was inoculated into various flask containing 20 ml of starch broth, with respective starch concentration, NaCl concentrations and pH value separately. All the flasks were incubated at 30°C for 7-8 days. Then the cell free filtrates of the culture were collected from all the media separately and were tested against the actively growing sensitive culture of *Serratia marcescens* by agar well diffusion method. The plates were incubated at 37° C for 24 hours followed by the measurement of zone of inhibition. The effect of starch as a carbon source on the growth and antimicrobial compound production was checked by using starch broth supplemented with various concentrations of starch (0, 0.5, 1 and 1.5%), 10% NaCl. The effect of the salt concentration on growth and the antimicrobial compound production was studied by using 1% starch broth provided with varying salt concentrations (0, 5, 10 and 15% w/v NaCl). The effect of pH on growth and antimicrobial compound production was studied by using 1% starch broth supplemented with 10% w/v NaCl and the range of pH (8, 9, 10 and 11).

### Extraction of Antimicrobial Metabolites

The strain *Nocardiopsis_sp. Al-H10-1* was cultivated in starch broth with 10% NaCl on a rotary shaker (120 rpm) at 37 °C for 8 days. The cell-free extract was obtained by filtration of broth culture using Whatman No. 1 filter paper. Equal volume of ethyl acetate was added to the culture filtrate for the extraction of the bioactive compounds. The mixture was added to 250 ml glass flask, sealed with cotton plug, followed by aluminum foil to reduce evaporation of the organic solvent and placed on shaker for 2 hours at 150 rpm. Post agitation the mixture was transferred to a separating funnel to generate different layers; the organic layer that contained the secondary metabolites and the aqueous layer. The crude extract was obtained by concentrating the solvent by evaporation and stored at 4°C for further use.

### Identification of Antimicrobial Metabolites by Using Gas Chromatography-Mass Spectrometry (GC-MS) Analysis

Identification of the chemical compounds present in the crude ethyl acetate extract of *Nocardiopsis_sp. Al-H10-1* was carried out by Gas Chromatography-Mass Spectrometry (GC-MS) analysis. Analysis was conducted on a capillary column (Rxi-5ms, 30m, 0.25 mm id, 0.25 µm film thickness) with the following conditions: constant flow of Helium, 1.0 ml min^-^1; the fixed inlet temperature, 285°C throughout the analysis; injection volume, 3 µl in the linear with an open purge valve (30:1 split ratio); Linear velocity: 36.8 cm/sec; Pressure: 65.0 kPa; Purge flow: 3.5 ml/min; Column flow: 1 ml/min; Oven ramp: 80 °C holds for 2.0 min, 18 °C/min to 260 °C holds for 6.0 min, 4°C/min to 285 °C holds for 6.0 min; Total run time: 30.25 min. The MS instrument with Ion source temperature: 200 °C; Interface temperature: 300 °C; Solvent cut time: 5.0 min; Detector voltage: 1 kV; Acquisition mode: Scan mode; Scan speed: 909; Event time: 0.78 sec; Starting m/z: 40 to 700 m/z. The peaks were identified by comparing the mass spectra with the National Institute of Standards and Technology (NIST, USA) library.

## RESULTS

### Isolation and Morphology of the Organism

*Nocardiopsis_sp. Al-H10-1*, a halophilic actinomycete was isolated from the site 20m away from Alang sea shore, Gulf of Khambhat, Western India. The isolate was characterized on the basis of its cell and colony morphology and Gram ‘s reaction. The colonies were medium sized, irregular shaped, irregular, slightly raised, rough, and opaque with creamy white pigmentation. It was Gram-positive, having a filamentous, long thread-like structure. It started sporulation on starch casein agar after 3 days of incubation with a fluffy mass of spores.

### Biochemical Characteristics and Antibiotic Susceptibility Test

*Nocardiopsis_sp. Al-H10-1* is a Gram-positive, filamentous and sporulating organism. Various biochemical tests were performed for primary characterization of the isolate. *Nocardiopsis_sp. Al-H10-1* showed positive results for Voges-Proskauer test, Ammonia production, Catalase test, Citrate utilization, Triple sugar iron test, Starch hydrolysis, Casein hydrolysis and Gelatin hydrolysis while organism showed negative results for Methyl red test, Indole production, Urea hydrolysis test, Nitrate reduction test and H_2_S production. The organism was grown in broth, supplemented with various carbohydrates like Mannitol, Sucrose, Xylose, Glucose, Maltose and Lactose to check its ability to utilize various carbohydrates and their fermentation pattern. Among the six provided carbohydrates, the organism was able to ferment glucose, sucrose, and maltose but not mannitol, xylose and lactose. The results of antibiotic susceptibility test showed that the isolate showed the highest sensitivity towards Rifampicin followed by Vancomycin, Ciprofloxacin, Gentamicin, Chloramphenicol, Streptomycin, Tetracycline, Kanamycin, Penicillin and Polymyxin-13 while the isolate was resistant towards Nalidixic acid, Ampicillin and Methicillin (Table 1).

**Table 1:** Biochemical characteristics of *Nocardiopsis_sp. Al-H10-1*

| Biochemical Test | Result |
| --- | --- |
| Methyl red test | - |
| Voges-Proskauer test | + |
| Indole production | - |
| Urease test | - |
| Nitrate reduction test | - |
| Ammonia production test | + |
| Catalase test | + |
| H<sub>2</sub>S production test | - |
| Citrate utilization test | + |
| TSI | + |
| Gelatin hydrolysis test | + |
| Casein hydrolysis test | + |
| Starch hydrolysis test | + |
| **Utilization of Carbohydrates** |  |
| Mannitol | - |
| Sucrose | + |
| Xylose | - |
| Glucose | + |
| Maltose | + |
| Lactose | - |
| **Antibiotic Susceptibility Test** |  |
| **Antibiotic (Concentration)** | **Zone of Inhibition (mm)** |
| Kanamycin (30 µg) | 7 (S) |
| Streptomycin (10 µg) | 21 (S) |
| Nalidixic acid (30 µg) | 0 (R) |
| Gentamicin (10 µg) | 22 (S) |
| Rifampicin (5 µg) | 32 (S) |
| Ciprofloxacin (5 µg) | 23 (S) |
| Ampicillin (10 µg) | 0 (R) |
| Polymyxin-13 (300 units/disc) | 1 (S) |
| Tetracycline (30 µg) | 18 (S) |
| Chloramphenicol (30 µg) | 22 (S) |
| Vancomycin (30 µg) | 28 (S) |
| Methicillin (5 µg) | 0 (R) |
| Penicillin (10 units/disc) | 5 (S) |

### Molecular Identification of Nocardiopsis_sp. Al-H10-1 through 16S rRNA Sequencing

The 16S rRNA gene sequencing of the strain *Nocardiopsis_sp. Al-H10-1* showed the presence of 1457 bp long 16S rRNA gene in the genomic sequence. The sequence was submitted to NCBI, GenBank, Maryland, USA with accession number (KF384482). The phylogenetic tree was constructed using neighbor-joining with Kimura 2-state parameter and pairwise-deletion model analysis implemented in the program MEGA software version X (Kumar S et. al., 2018) and also evaluated further in a bootstrap analysis of 1,000 replicates (Figure 1). The molecular characterization through 16S rRNA gene sequencing revealed the organism belonging to *Nocardiopsis sp*.

**Fig 1.**
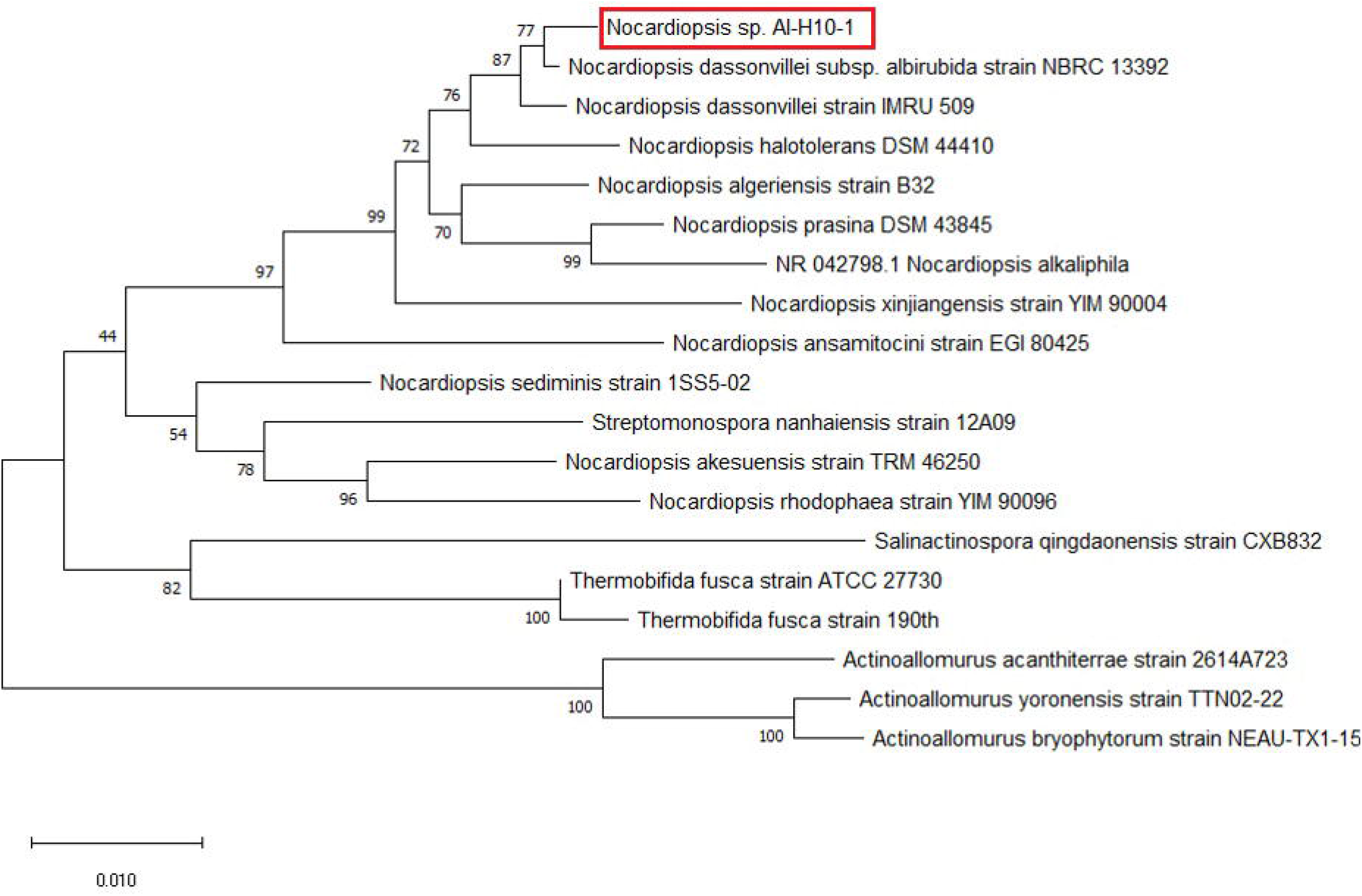
Neighbor-joining tree of the strains and representative species of the genus *Nocardiopsis and Streptomyces* based on nearly complete 16S rRNA gene sequences. Numbers at the nodes indicate the levels of bootstrap support based on 1000 resampled data sets. The scale bar indicates 0.01substitution per nucleotide position.

### The Antimicrobial Potential

The antimicrobial potential of the isolate was checked against six Gram-positive, eight Gram-negative bacteria and three fungi. The isolate *Nocardiopsis_sp. Al-H10-1* inhibited the growth of the total five test organisms including four Gram negative bacteria *Pseudomonas aeruginosa, Serratia marcescens, Enterobacter aerogenes, Klebsiella pneumonia* and one Gram positive bacterium *Bacillus megaterium* was inhibited (Figure 2). The highest inhibition was recorded against *Serratia marcescens*, however the isolate didn ‘t show any inhibitory effect against any of the fungi tested for the same.

**Fig 2.**
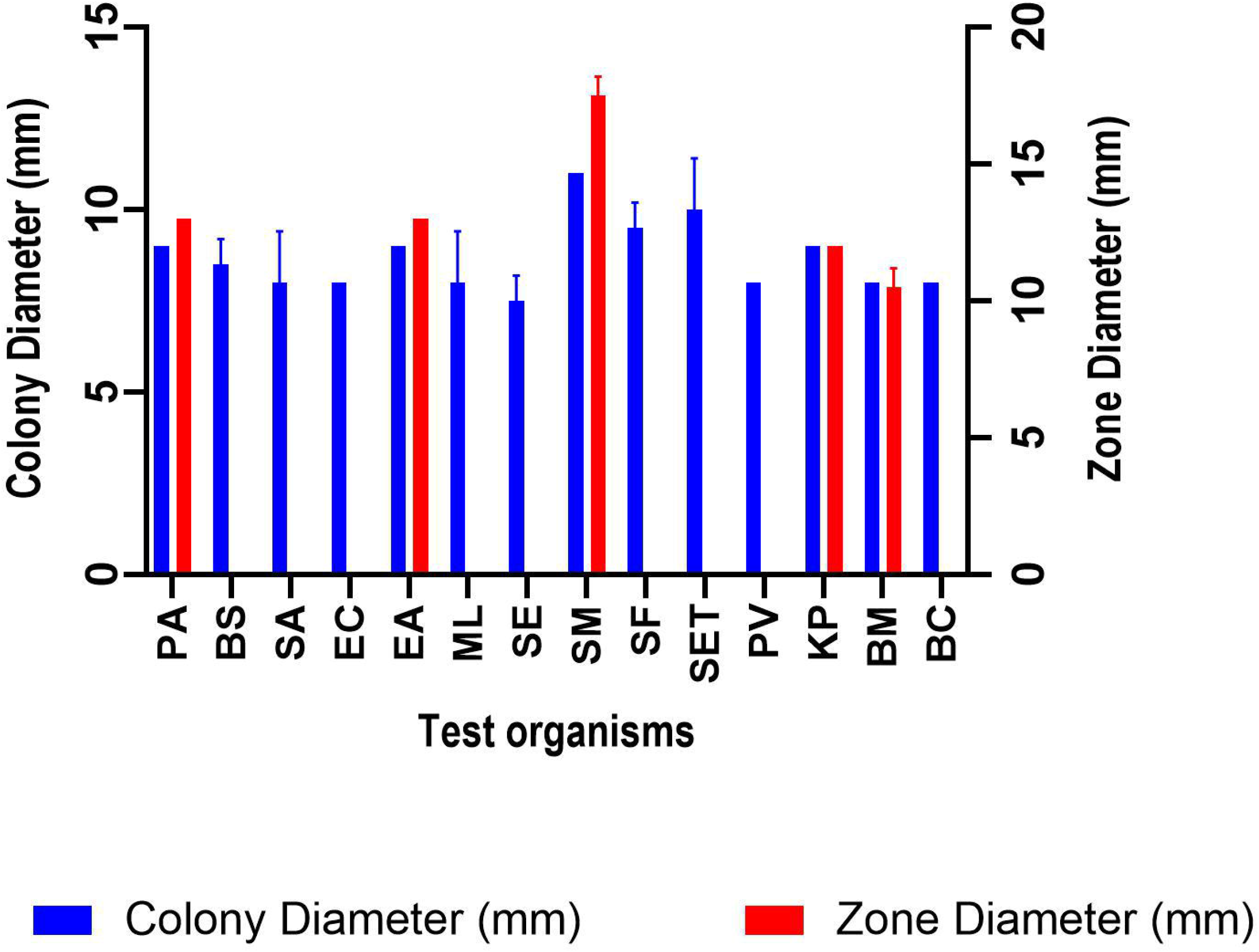
Antibacterial activity of *Nocardiopsis_sp. Al-H10-1* isolate against various test organisms; blue bar (Colony diameter, mm), Green bar (Zone diameter, mm)

### Optimization of Growth Conditions for Antimicrobial Compound Production

To achieve the highest production of antimicrobial compounds, effect of various growth conditions were studied. Optimization of various growth conditions showed that the highest antimicrobial compound production was obtained in the presence of 0.5 % starch, 10% NaCl and pH 9 (Figure 3-5).

**Fig 3.**
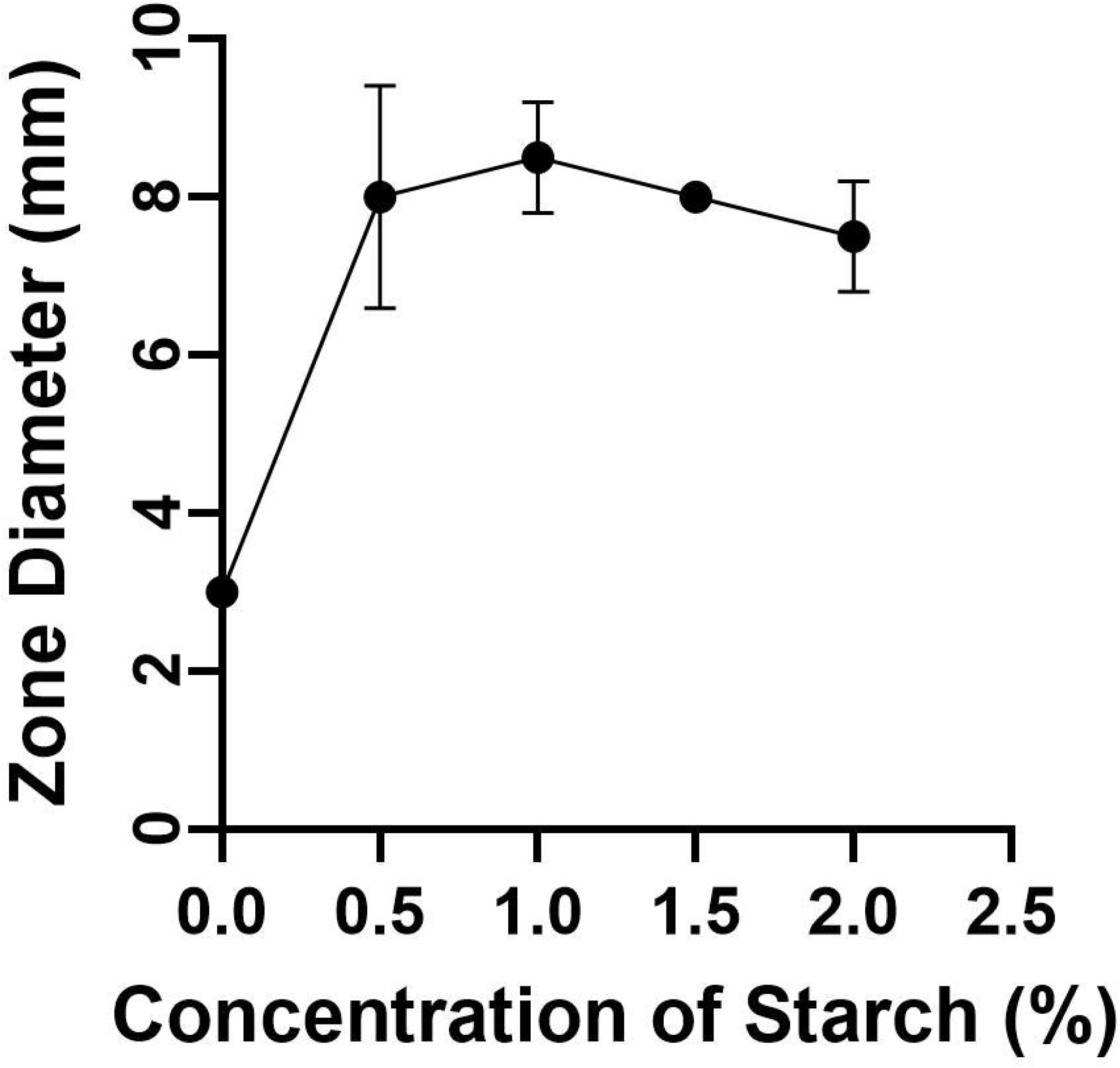
Effect of starch concentrations on growth and antimicrobial compounds production by *Nocardiopsis_sp. Al-H10-1* against *Serratia marcescens*

**Fig 4.**
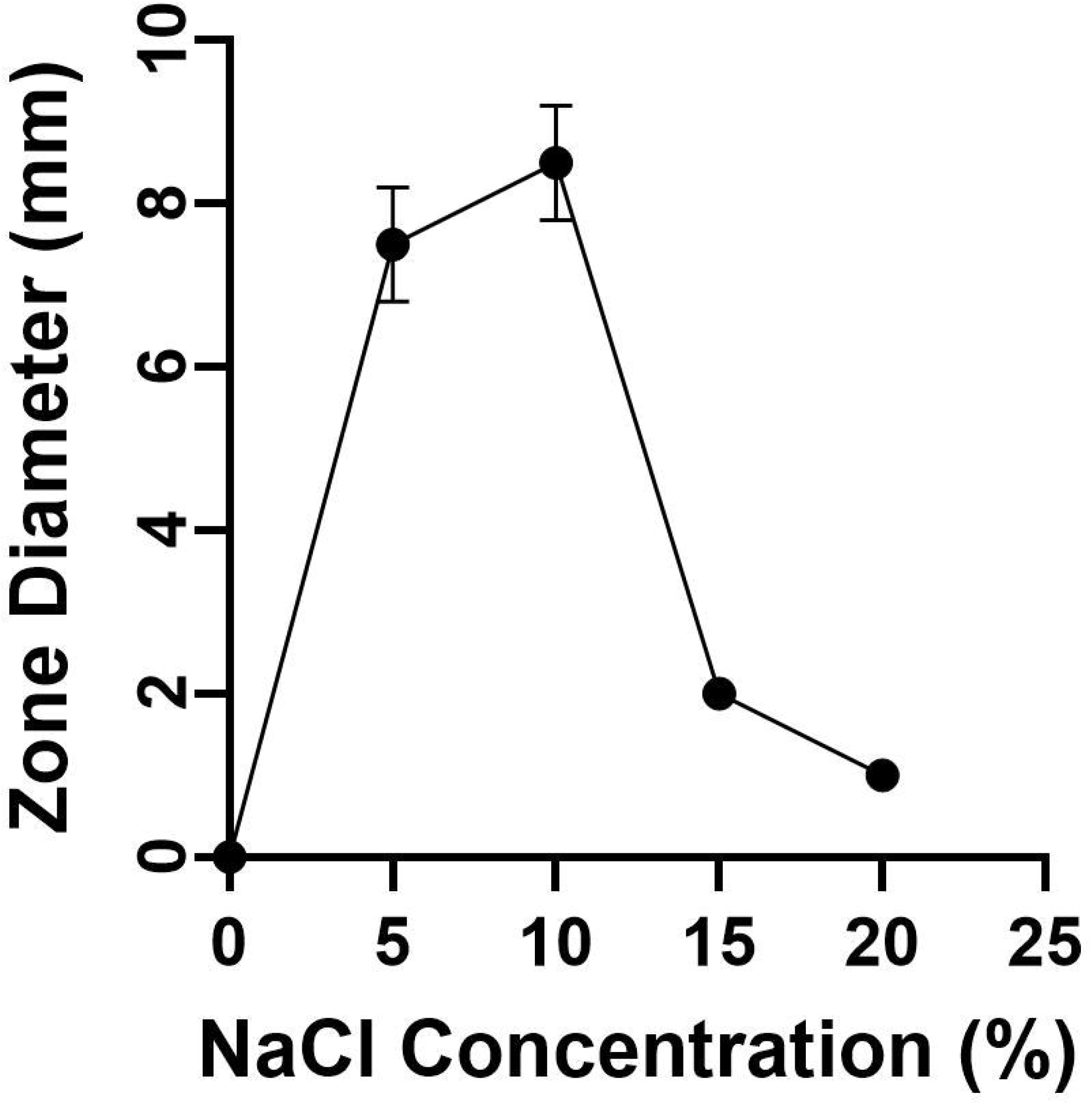
Effect of salt concentrations on growth and antimicrobial compounds production by *Nocardiopsis_sp. Al-H10-1* against *Serratia marcescens*

**Fig 5.**
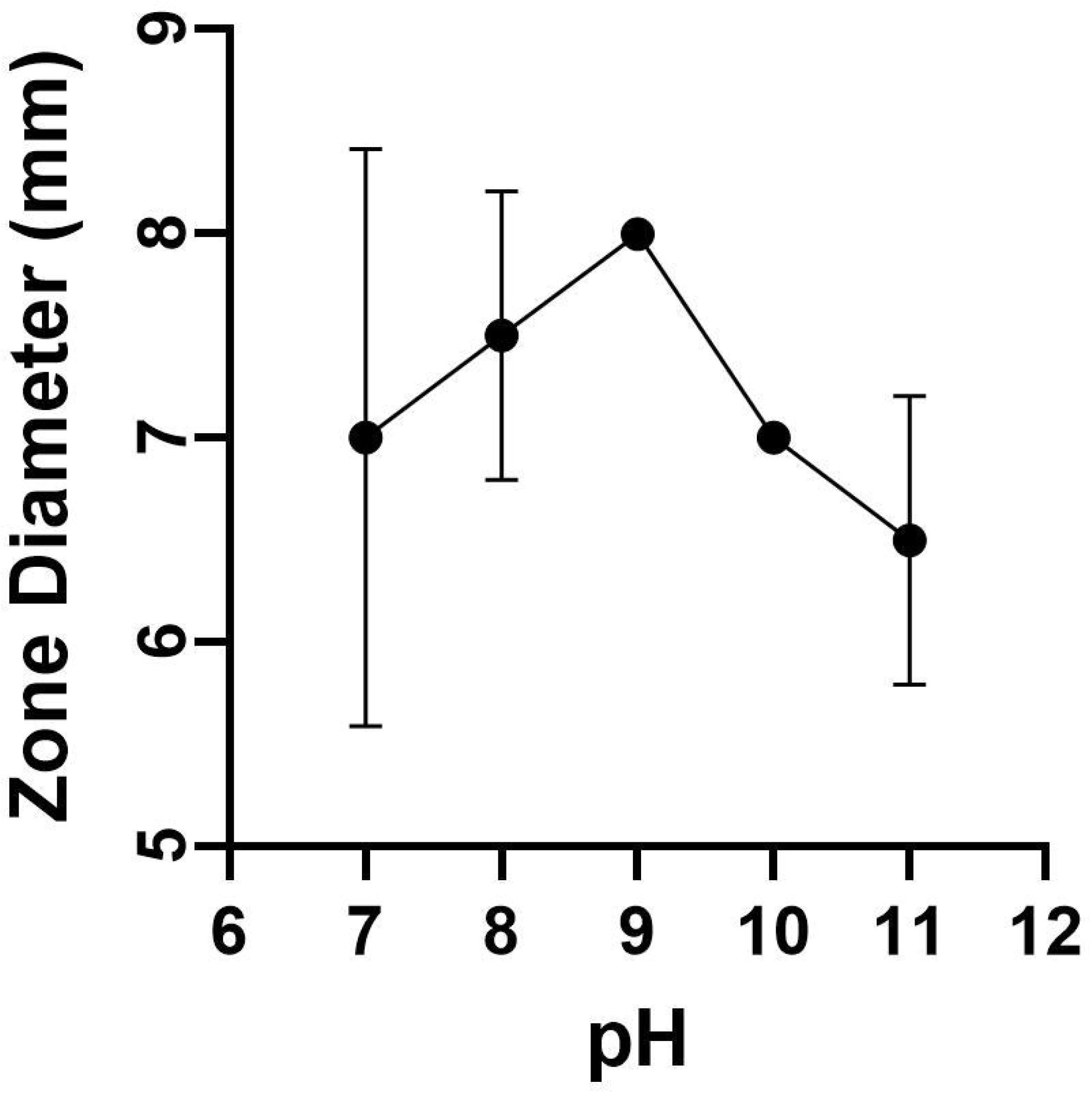
Effect of pH on growth and antimicrobial compounds production by *Nocardiopsis_sp. Al-H10-1* against *Serratia marcescens*

### Identification of Bioactive Compounds by GC-MS Analysis of Ethyl Acetate Extract

Identification of the bioactive compounds, present in ethyl acetate extract, was carried out using GC-MS analysis. The GC-MS chromatogram of the *Nocardiopsis_sp. Al-H10-1* crude extract showed a total of 17 peaks (Figure 6). When compared with the NIST database, the nearest compound hits for those peaks were found (Table 2).

**Table 2:** GC-MS analysis of *Nocardiopsis_sp. Al-H10-1* ethyl acetate extract

| Peak No. | Retention Time | Name of the Compound | Molecular Formula | Molecular Weight | Peak Area % | Activity | Reference |
| --- | --- | --- | --- | --- | --- | --- | --- |
| 1 | 8.045 | Dodecane, 2-methyl- | C<sub>13</sub>H<sub>28</sub> | 184.36 | 2.42 |  |  |
| 2 | 8.290 | Tetradecane | C<sub>14</sub>H<sub>30</sub> | 198.39 | 2.99 | Antimicrobial | Girija S et. al., 2014 |
| 3 | 8.471 | 2,4-bis(1,1-dimethylethyl)- Phenol | C<sub>14</sub>H<sub>22</sub>O | 206.32 | 3.84 | Antibacterial | Nandhini SU et. al., 2015;<br>Yogeswari S et. al., 2012 |
| 4 | 8.602 | Pentadecane | C<sub>15</sub>H<sub>32</sub> | 212.41 | 2.29 | Antimicrobial | Girija S et. al., 2014 |
| 5 | 9.714 | Pentadecane | C<sub>15</sub>H<sub>32</sub> | 212.41 | 3.37 | Antimicrobial | Girija S et. al., 2014 |
| 6 | 9.978 | Heptadecane | C<sub>17</sub>H<sub>36</sub> | 240.5 | 2.51 | Antimicrobial | Rahbar N et. al., 2012 |
| 7 | 10.243 | Heptadecyl heptafluorobutyrate | C<sub>21</sub>H<sub>35</sub>F<sub>7</sub>O<sub>2</sub> | 452.5 | 2.51 |  |  |
| 8 | 10.778 | Octadecane | C<sub>18</sub>H<sub>38</sub> | 254.5 | 3.61 | Antibacterial | Nazemi M et. al., 2010 |
| 9 | 10.979 | Heptadec-1-ene | C<sub>17</sub>H<sub>34</sub> | 238.5 | 3.09 |  |  |
| 10 | 11.149 | Benzenepropanoic acid, 3,5-bis(1,1-dimethylethyl-4-hydroxy-, methyl ester | C<sub>17</sub>H<sub>26</sub>O<sub>3</sub> | 278.4 | 2.15 |  |  |
| 11 | 11.260 | Dibutyl phthalate | C<sub>16</sub>H<sub>22</sub>O<sub>4</sub> | 278.34 | 25.69 | Antimicrobial | Roy RN et. al., 2006; Zhang H et. al., 2018 |
| 12 | 11.901 | Heneicosane | C<sub>21</sub>H<sub>44</sub> | 296.6 | 2.31 | Antimicrobial | Nandhini SU et. al., 2015 |
| 13 | 12.109 | 8-methyl-Heptadecane | C<sub>18</sub>H<sub>38</sub> | 254.5 | 1.48 |  |  |
| 14 | 12.407 | Pentatriacontane | C<sub>35</sub>H<sub>72</sub> | 492.9 | 2.62 | Antibacterial | Nithya TG et. al., 1972 |
| 15 | 18.716 | Squalene | C<sub>30</sub>H<sub>50</sub> | 410.7 | 1.43 | Antibacterial | Sermakkani M and Thangapandian V, 2012 |
| 16 | 28.569 | Diethyl-heptoxy-octadecoxysilane | C<sub>29</sub>H<sub>62</sub>O<sub>2</sub>Si | 470.9 | 28.07 |  |  |
| 17 | 31.916 | Pentacyclo[19.3.1.1(3,7).1(9,13).1(15,19)]octa | C<sub>40</sub>H<sub>48</sub>O<sub>4</sub> | 592.8 | 9.60 |  |  |

**Fig 6.**
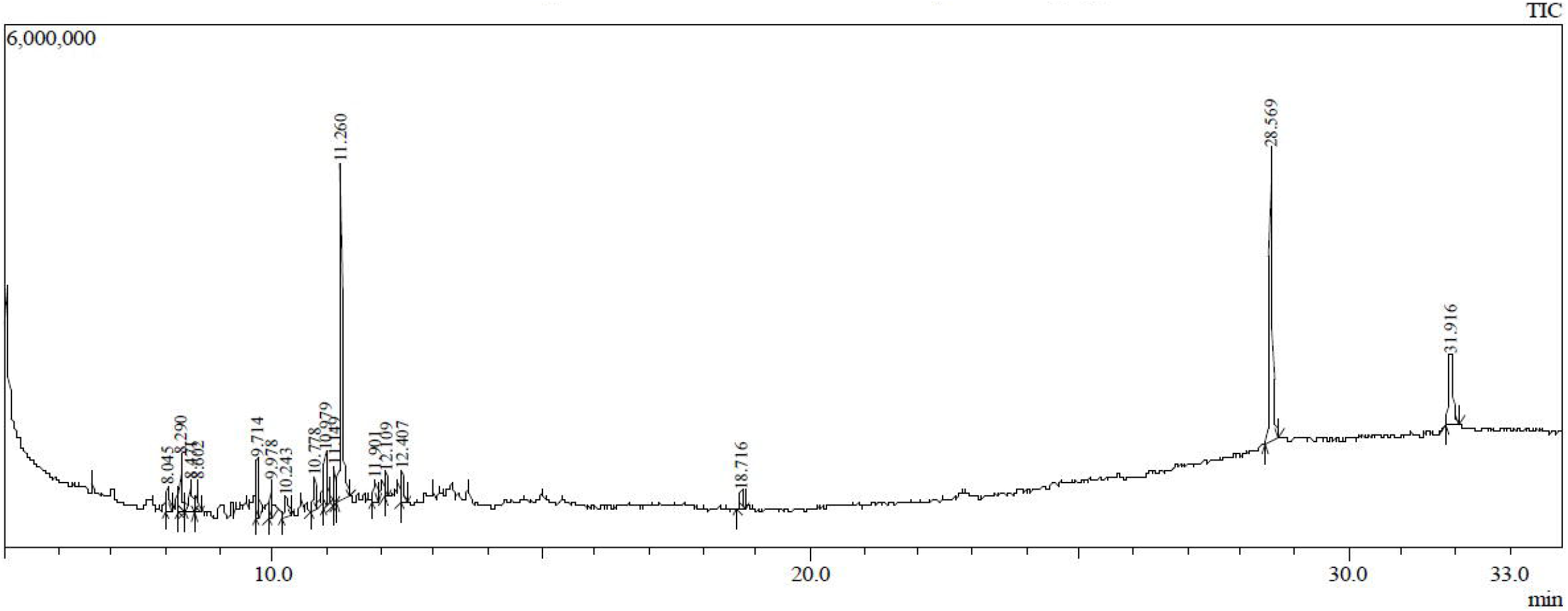
GC-MS chromatogram of *Nocardiopsis_sp. Al-H10-1* ethyl acetate extract

## DISCUSSION

From 2007 to 2017 (the duration of 10 years), total 177 new species of rare actinomycetes were reported from the marine habitats representing 33 families from which 3 were novel families and 29 novel genera (Subramani R and Sipkema D, 2019). The study of Berdy J, 2012 suggested that there are approximately 500,000 naturally occurring biological compounds, from which approximately 70,000 are microbially-derived with 29% is exclusively derived from actinomycetes. Marine actinomycetes are the prominent source of enzymes which can be produced on a large scale and can be used in various industries, such as proteases, cellulases, chitinases, amylases, xylanases, and other enzymes (Vaijayanthi G et. al., 2016). They are also known as the most valuable source of naturally occurring novel antimicrobial compounds (Fiedler Forsyth MP et. al., 1971; Hassanshahian M and Mohamadian J, 2011). So we focused on halophilic actinomycetes for the production of novel antimicrobial compounds in our research. *Nocardiopsis_sp. Al-H10-1* was successfully isolated from the marine water sample from Alang sea shore using selective media supplemented with 10% NaCl. On the basis of morphological and molecular characterization, the isolate was identified as Gram-positive, filamentous, long thread-like structure containing *Nocardiopsis_sp. Al-H10-1* able to inhibit the growth of four Gram negative bacteria including *Pseudomonas aeruginosa, Serratia marcescens, Enterobacter aerogenes, Klebsiella pneumonia* and one Gram positive bacterium *Bacillus megaterium*. Das R et. al. (2018) isolated total 172 actinobacteria using soil samples collected from two microbiologically unexplored forest ecosystems, located in the Eastern Himalayan Biodiversity hotspot region i.e. Nameri National Park (NNP) and Panidehing Wildlife Sanctuary (PWS) by using the same isolation techniques used by us, they further screened 24 strong antimicrobial activity showing isolates using the same spot inoculation method. Biochemical tests of *Nocardiopsis_sp. Al-H10-1* showed that the isolate is able to utilize various carbon sources like glucose, citrate, starch, sucrose and maltose; and nitrogen sources like casein and gelatin with production of ammonia. The organism is also able to produce various industrially important enzymes including amylase, proteases like caseinase and gelatinase. The isolate was able to produce catalase which indicates the organism is aerobic in nature. The organism was able to ferment various carbohydrates like glucose, sucrose, and maltose. The negative results for H_2_S production and nitrate reduction test indicated that the isolate is not able to utilize more complex protein contents and sulfur containing amino acids from the surrounding habitats. Antibiotic sensitivity/resistant profiling is widely used to study microbial diversity in combination with biochemical and molecular characterization of the isolates (Litzner BR et. al., 2006). The antibiotic sensitivity/resistant profiling showed that the isolate is sensitive towards range of protein synthesis inhibiting antibiotics such as chloramphenicol, gentamycin, streptomycin and tetracycline. The highest sensitivity showed towards RNA synthesis inhibiting rifampicin followed by cell wall synthesis inhibiting vancomycin while the isolate showed resistance towards DNA synthesis inhibitor nalidixic acid and cell wall synthesis inhibitors ampicillin and methicillin. The phylogenetic analysis of the isolate showed that *Nocardiopsis_sp. Al-H10-1* showed the highest similarity with *Nocardiopsis halotolerans* DSM44410 strain NBRC100347 (98.63%) and the lowest similarity with *Actinoallomurus acanthiterrae* strain 2614A723 (91.59%). The antimicrobial compounds produced by *Nocardiopsis_sp. Al-H10-1* were extracted by using ethyl acetate (1:1 v/v) from filtrate broth. Charousová I et. al., 2019 also extracted the antimicrobial compounds produced by selected actinomycetes isolated using aridic soil sample collected from Karoo, South Africa. Identification of the bioactive compounds, present in ethyl acetate extract, was carried out using GC-MS analysis. The GC-MS chromatogram of the *Nocardiopsis_sp. Al-H10-1* crude extract showed a total of 17 peaks. When compared with the NIST database, the nearest compound hits for those peaks were found. GC-MS analysis indicated the presence of 16 compounds including alkanes (Tetradecane, Pentadecane, Heptadecane, Octadecane, Henicosane, Pentatriacontane); alkenes (Heptadec-1-ene); triterpene (Squalene); phenol (2,4-bis(1,1-dimethylethyl)-Phenol); Dibutyl phthalate; Dodecane-2-methyl-; Heptadecyl heptafluorobutyrate; Benzenepropanoic acid, 3,5-bis(1,1-dimethylethyl-4-hydroxy-, methyl ester; 8-methyl-heptadecane; Diethyl-heptoxy-octadecoxysilane; Pentacyclo[19.3.1.1(3,7).1(9,13).1(15,19)]octa. From 16 identified compounds, 10 compounds have already been reported to have antimicrobial activity (Girija S et. al., 2014; Nandhini SU et. al., 2015; Yogeswari S et. al., 2012; Rahbar N et. al., 2012; Nazemi M et. al., 2010; Roy RN et. al., 2006; Zhang H et. al., 2018; Nithya TG et. al., 1972; Sermakkani M and Thangapandian V, 2012). El-Naggar NE et. al., 2017 also reported presence of alkanes in the ethyl acetate extract of *Streptomyces anulatus* NEAE-94 using GC-MS analysis.

## ACKNOWLEDGEMENT

Financial Assistance from the Department of Biotechnology (New Delhi, India) is acknowledged.

